# Conspecifics, not plant reproductive tissues, reduce omnivore prey consumption

**DOI:** 10.1101/596015

**Authors:** S. Rinehart, J.D. Long

## Abstract

Plant reproductive tissues (PRTs) can decrease (via reduced consumption) or increase (via numerical response) an omnivores consumption of animal prey. Although PRTs can increase predation pressure through numerical responses of omnivores, PRTs may also suppress predation by increasing omnivore interactions with conspecifics. Despite this potential, studies of the impacts of PRTs on predation by omnivores often overlook the effect of these tissues on intraspecific interactions between omnivores. We designed three studies to examine how PRTs and conspecific density impact prey consumption by ladybeetle omnivores. First, we assessed how PRTs impact scale insect consumption by isolated ladybeetles. Second, we measured how PRTs influence ladybeetle prey suppression when numerical responses were possible. Third, because initial experiments suggested the consumption rates of individual ladybeetles depended upon conspecific density, we compared per capita consumption rates of ladybeetles across ladybeetle density. PRTs did not influence prey consumption by isolated ladybeetles. When numerical responses were possible, PRTs did not influence total predation on prey despite increasing ladybeetle density, suggesting that PRTs decreased per capita prey consumption by ladybeetles. The discrepancy between our lab and field studies is likely a consequence of differences in ladybeetle density - the presence of only two other conspecifics decreased per capita prey consumption by 76%. Our findings suggest that PRTs may not alter the population level effects of omnivores on prey when omnivore numerical responses are offset by reductions in per capita predation rate.

## Introduction

Omnivory (i.e. consuming resources from multiple trophic levels) [1] is ubiquitous within several taxa (e.g. birds, mammals, reptiles, insects, and fishes) and influences the structure and function of communities [2-3]. Interactions between omnivores and their plant and animal prey can account for up to 78% of species’ links in food webs [4]. Despite their prevalence, we lack a basic understanding about how plant reproductive tissues (hereafter, PRTs) affect interactions between omnivores and their prey in natural systems. Some studies suggest that PRTs decrease prey consumption by omnivores [5-6], whereas others suggest the opposite [7]. This discrepancy may be exacerbated by methodological approaches and the spatial scale of the study [8-9]. For instance, many omnivory studies focus on isolated omnivores feeding on a sub-set of possible resources, which only allows omnivore consumption to depend on resource density and the availability of PRTs [8,10]. Such approaches fail to allow important intraspecific interactions (e.g., mating, cannibalism, and competition) and interspecific interactions (e.g., predation and competition), whose occurrence may be altered by PRTs [9, 11-13]. Understanding how PRTs and conspecific density affect prey consumption by omnivores may help predict when and where omnivores exert top-down control on prey populations.

PRTs may suppress omnivore consumption of prey if these resources are equally (or more) palatable than prey or provide critical habitat structure [14-16]. For example, when PRTs (pollen) was available, omnivorous phytoseiid mites (*Iphiseius degenerans*) consume fewer prey (larval *Euseius stipulatus*), leading to lower prey mortality [13]. Similarly, omnivorous big-eyed bugs (*Geocoris punctipes*) consume fewer pea aphids (*Acyrthosiphon pisum*) and are less effective at regulating pea aphid populations when high quality plant tissue (lima bean pods) is locally available [5].

In contrast, omnivore prey consumption may increase in the presence of PRTs if these resources increase the local abundance of omnivores through a numerical response (i.e. aggregation and enhanced fitness) [5, 15-18] or by lengthening omnivore persistence in habitats with low prey densities [1, 17, 19-21]. For instance, habitat patches containing high densities of PRTs (lima bean pods) had larger populations of omnivores and omnivores were less-likely to emigrate from patches containing lima bean pods [17].

Elevated conspecific density, due to omnivore numerical responses to PRTs, may increase the frequency of intraspecific interactions (i.e. interactions between conspecifics such as mating, cannibalism, territoriality, competition) [22-23], thereby decreasing the per capita prey consumption by omnivores [24-26]. For example, flatworm predators (*Stenostomum virginanum*) reduce their per capita predation rates on protozoan prey in the presence of conspecifics [26]. Similarly, larval tiger salamanders lower their foraging rates when larger conspecifics are present, likely to minimize their risk of being cannibalized [27]. While the effects of PRTs on intraspecific (e.g., cannibalism) and intraguild interactions have been well-studied [see 9, 11-13], few studies have aimed to understand how changing omnivore densities (associated with numerical responses to PRTs) can indirectly affect prey population dynamics in the field.

Most empirical studies testing the impacts of PRTs on omnivore prey consumption have used insect omnivores as model systems [see 5, 7, 17, 28-32]. Many insects, like omnivores belonging to other taxa like foxes, sharks, and coyotes, are active predators [33-37], meaning that they continuously search for prey [38]. This suggests that studies of omnivorous insects may inform how non-insect omnivores impact prey populations.

Here, we assessed how PRTs and conspecific density affect the foraging behavior of an omnivorous salt marsh ladybeetle (*Naemia seriata*) feeding on scale insects (*Haliaspis spartinae*). We used laboratory mesocosms to assess the impacts of PRTs [i.e. cordgrass (*Spartina foliosa*) flowers] on ladybeetle per capita consumption of scale insects. We paired laboratory mesocosms with a field study to assess the impact of cordgrass flowers on ladybeetle and scale insect density under natural conditions, where numerical responses were possible. Finally, to reconcile our laboratory and field studies, we conducted a laboratory no-choice feeding assay to assess how conspecific density impacts ladybeetle per capita scale insect consumption.

## Methods

### Study system

We assessed how PRTs and conspecific density influence the ability of the omnivorous ladybeetle, *Naemia seriata* (hereafter ladybeetle), to suppress populations of its insect prey, the armored scale insect *Haliaspis spartinae* (hereafter, scale insects). Scale insects are specialist phloem-feeders on the foundational salt marsh plant, *Spartina foliosa* (hereafter, cordgrass). We used this ladybeetle-scale insect model system for three reasons. First, ladybeetles in this system are facultative omnivores, as access to cordgrass pollen facilitates ladybeetle survival in the absence of other dietary resources [18]. Specifically, adult ladybeetles provided only access to cordgrass pollen survived 1.97-times longer than ladybeetles provided access to no food resources. This suggests that in the absence of other prey resources, adult ladybeetles likely consume cordgrass pollen to promote their longevity. Second, adult ladybeetles show resource-dependent aggregation in the field, with ladybeetles tending to preferentially aggregate to habitats containing both scale insects and cordgrass flowers over habitats lacking these resources [18]. Third, adult ladybeetles often aggregate with conspecifics on cordgrass flowers (S. Rinehart and J.D. Long *unpublished data*), suggesting that cordgrass flowers may be a hub of ladybeetle intraspecific interactions (e.g. mating and territoriality).

### Effect of cordgrass flowers on scale insect consumption by isolated ladybeetles

To test how Flower Access [2 Levels: Flower Access Present (FA+), Flower Access Absent (FA-)] affects consumption of scale insects by individual adult ladybeetles, we conducted a mesocosm experiment at the San Diego State University Coastal and Marine Institute Laboratory (CMIL). On 20-July-2015, we collected 20 sediment plugs (15 x 15 cm; diameter x deep) each containing a single flowering cordgrass stem infested with scale insects from Sweetwater Marsh (South San Diego Bay; 32° 38’ 15.8’’N, 117° 06’ 37.5’’W). We observed pollen on all cordgrass stems collected at this time. We planted cordgrass stems and field-collected sediment in 2.6 L plant pots with holes for drainage (Elite Nursery Containers; 300 Series). We used toothbrushes to remove all non-scale insect resources (e.g., leafhoppers of pollen) from cordgrass leaves and to standardized initial mean total scale insect density to 559 ± 73 insects stem^-1^ (mean ± SE). We collected ladybeetles from two sites, Sweetwater Marsh and San Dieguito Lagoon (32° 58’ 40.4’’N, 117° 14’ 32.8’’W).

We placed all potted plants in an outdoor, flow-through seawater table. Plants were rearranged randomly each week. We connected our seawater table to a tidal control system that automatically changed tank tidal conditions [between high (plant pots submerged) and low (plant pots not submerged)] at preset intervals creating tidal conditions like those experienced by cordgrass at Sweetwater Marsh at a tidal height of 1.5 m above sea-level. We let potted plants acclimate to tank conditions for two weeks prior to the experiment.

On 03-Aug-2015, we randomly assigned potted cordgrass plants to a Ladybeetle (Present, Absent) and a Flower Access treatment [Flower Access Present (FA+), Flower Access Absent (FA-)]. All treatments had scale insects present (n=5). In the Ladybeetle Present treatment, we introduced a single adult ladybeetle into each replicate. We replaced ladybeetles every other week, as we experienced a 10% mortality rate each week. In FA-treatments, we placed cordgrass flowers in 16 x 14 cm Glad Fold-Top plastic bags (The Glad Company; Oakland, California). We secured bags to plants with a cable tie. These bags prevented ladybeetle access to the flowers and thus indirectly manipulated their ability to access cordgrass pollen. In FA+ treatments, we did not restrict ladybeetle access to cordgrass flowers and pollen. However, we controlled for the cable tie by attaching a cable tie to all cordgrass stems in FA+ treatments. We prevented insect dispersal among replicates by covering each entire replicate with nylon insect mesh (54 x 50 cm, height x width, mesh size = 1 mm). We maintained this experiment for 6 weeks until 14-Sept-2015.

To assess the effect of Flower Access and Ladybeetles on scale insect density, we monitored adult and juvenile (hereafter “crawler”) scale insect density every two weeks. Adult and crawler scale insects can be distinguished by their mobility and morphology (e.g. Unlike crawlers, adults are immobile and produce a white waxy test). We corrected for natural fluctuations in scale insect density by pairing replicates from the Ladybeetle Present and Ladybeetle Absent treatments and using the formula: P_i_ (A_f_/ A_i_) – P_f_ [39]. Here, P_i_ and P_f_ represent the initial and final scale insect density of Ladybeetle Present treatments and A_i_ and A_f_ represent the initial and final scale insect density of Ladybeetle Absent treatments. This correction allowed us to detangle natural variation in scale insect population dynamics from effects of Flower Access. We then compared our corrected adult, crawler, and total scale insect per capita consumption by ladybeetles between Flower Access (FA+, FA-) treatments using a series of two-sample t-tests. All corrected scale insect consumption data were square-root transformed. We conducted all statistical analyses in JMP v. 13 (www.jmp.com).

### Effect of cordgrass flowers on ladybeetle aggregation and scale insect consumption

To assess how Flower Access [(Flower Access Present (FA+), Flower Access Absent (FA-)] influences ladybeetle aggregation and consumption of scale insects, we conducted a fully-factorial study at San Dieguito Lagoon. On 18-Aug-2016, we established 20 - 0.25m^2^ circular plots (separated by at least 1m) in a monospecific cordgrass stand infested with scale insects. Although ladybeetles may feed on other prey resources (e.g., leafhoppers), they likely constitute only a small portion of ladybeetle diets, as alternative prey resources are rare compared to scale insects. For example, at the experimental site, scale insect density was 16,177 ± 2,174 per 0.25m^2^ (mean ± SE), while leafhopper density was only 25 ± 2.8 per 0.25m^2^ (mean ± SE; S.A. Rinehart *unpublished data*). All plots started with at least four flowering cordgrass stems, a cordgrass stem density of 22 ± 1.1 (mean ± SE), and zero ladybeetle egg clutches. We randomly allocated plots to each treatment (n=10). In the FA-treatment, we covered all cordgrass flowers with 16 x 14 cm Glad Fold-Top plastic bags (The Glad Company; Oakland, California) and secured bags in place with a cable tie. In the FA+ treatment, we did not inhibit ladybeetle access to cordgrass flowers. However, we controlled for the presence of cable ties in the FA+ treatment by applying cable ties to all stems included in the study. We used plastic bags to inhibit ladybeetle access to cordgrass flowers rather than mesh bags, as plastic also inhibits the transmission of plant volatile cues [40].

To assess how Flower Access influences ladybeetle aggregation, we monitored the density of all ladybeetle life stages (adults, larvae, and egg clutches) in each plot weekly between 08-Aug-2016 and 22-Sept-2016. We determined the density of ladybeetle life stages using two-minute timed searches. During timed searches, we examined all stems in each plot, starting at the soil-air interface and working toward the apical meristem. All ladybeetle life stage densities were log transformed. We tested for effects of Flower Access on the density of each ladybeetle life stage using separate RM-ANOVAs with Flower Access as a fixed factor and week as the repeated measure.

To understand how Flower Access influences ladybeetle suppression of scale insect populations under field conditions, we recorded scale insect density on two focal cordgrass stems (all focal stems had flowers present) in each replicate on two dates (18-Aug-2016 and 22-Sept-2016). We summed the total scale insect density on focal stems in each plot for both timepoints and used this value to calculate the change in scale insect density per plot over the five-week study. The change in scale insect density was square-root transformed. We then compared the change in scale insect density (per two focal stems) between Flower Access (FA+ vs. FA-) treatments using a two-sample t-test.

### Effect of conspecific density on ladybeetle per capita scale insect consumption

Because 1) the impact of flower access on per capita consumption of scale insects differed between our laboratory and field studies (no effect in the laboratory, decreased consumption in the field) and 2) intraspecific interactions between ladybeetles also varied between these studies (absent in the laboratory, present in the field), we conducted a no-choice feeding assay to examine the influence of ladybeetle density on per capita consumption of scale insects. On 10-Nov-2017, we collected ladybeetles and flowering, scale-infested cordgrass stems (clipped at the air-soil interface) from San Dieguito Lagoon two hours prior to the study. Collected stems and ladybeetles were transported to the CMIL, where we counted the initial total scale insect density per cordgrass stem. Cordgrass stems were then randomly allocated to each Ladybeetle Density treatment: 0, 1, 2, or 3 per stem. We based the upper Ladybeetle Density treatment on survey data showing that adult ladybeetles tend to aggregate in groups of 3 ± 0.6 individuals per cordgrass stem. Sample size was five for all Ladybeetle Density treatments except the 0 treatment, which had three replicates. We then placed the clipped end of each cordgrass stem into its own 13 x 13 cm (height x diameter) cylindrical plastic container filled with 700 ml of tap water (to act as a vase) and enclosed the whole cordgrass stem and plastic container in a 54 x 13 cm (length x width) bag made with white nylon insect mesh (6 mm mesh opening). Finally, we introduced zero, one, two, or three adult ladybeetles to each replicate. We accidentally added four ladybeetles to one of the three ladybeetle treatments. All replicates were maintained at a mean temperature of 21.1°C with a 12:12 hour light-dark cycle (85.6 ± 5 μmol photons • m-2 • s -1 (PAR); Philips Natural Light 40W). After three days, we removed ladybeetles (no ladybeetles were lost or cannibalized during the study) and counted the final total scale insect density on all stems. We then calculated the total scale insects consumed (between all ladybeetles) and the per capita scale insect consumption of ladybeetles in all replicates.

Because there was no change in scale insect density in zero ladybeetle replicates during the study (one-sample t-test: t_2.00_ = 0.256, p = 0.589), we removed this treatment from further analysis and attributed all reductions in scale insect density to ladybeetle feeding. Using our 1,2, and 3 ladybeetle treatments, we tested for the effects of ladybeetle density (consistent through the study) on total scale insect consumption and the per capita consumption of ladybeetles using linear regressions with ladybeetle density as the independent factor. Total scale insects consumed (between all ladybeetles) and the per capita scale insect consumption were square-root transformed prior to analyses.

On the 2^nd^ and 3^rd^ days of the assay (11-Nov-2017 and 12-Nov-2017), we conducted behavioral observations of ladybeetles in all replicates. On each day, we recorded the location (i.e., plant leaf, plant stem, plant flower, or mesh bag) of each ladybeetle in each replicate between the hours of 08:00 and 10:00 am. We then calculated the number of ladybeetles in each replicate that were on any part of the plant (e.g., leaves, stem, or flower) at the time of observation. We tested for effects of ladybeetle density on the number of ladybeetles on any part of the plant using a RM-ANOVA with Ladybeetle Density as a fixed factor and Observation Day as the repeated measure.

## Results

### Effect of cordgrass flowers on scale insect consumption by isolated ladybeetles

Although ladybeetles have been observed consuming pollen (S.A. Rinehart *personal observation*), consumption of adult, crawler, and total scale insects by isolated ladybeetles was not affected by access to cordgrass flowers (Adults: t_6.59_ = 0.052, p = 0.96; Crawlers: t_5.32_ = 0.596, p = 0.576; Total: t_7.89_ = 0.216, p = 0.834; Fig 1).

**Fig 1.**
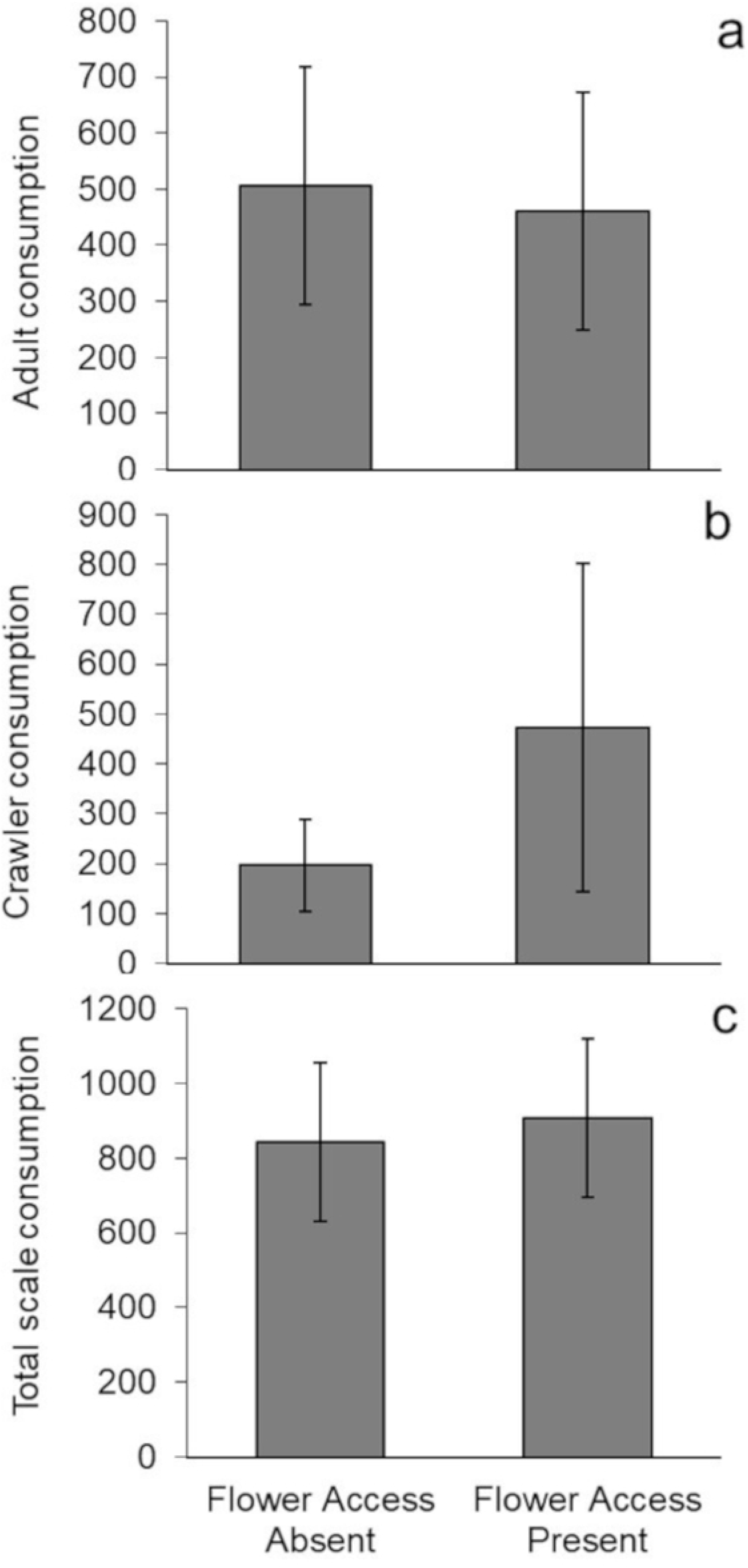
Effect of PRTs on isolated ladybeetle foraging. Control corrected mean (± SE) consumption by isolated adult ladybeetles of a) adult, b) crawler, and c) total scale insects (n = 5).

### Effect of cordgrass flowers on ladybeetle aggregation and scale insect consumption

In our field experiment, adult ladybeetle density depended on Flower Access (F_1,107_ = 43.69, p <0.001; S1 Table) and week (F_5,107_ = 6.73, p < 0.001). Ladybeetles increased with both factors. Flower Access and week also had an interactive effect on local adult ladybeetle density (F_5,107_ = 6.65, p < 0.001, Fig 2a). This interaction resulted from the differential effects of Flower Access on adult ladybeetle density through time. Specifically, adult ladybeetle density in plots with flower access increased by 412%, while adult ladybeetle density in plots without flower access actually decreased by 8% over the five-week study. Additionally, this effect was strengthened by differences in the initial adult ladybeetle density between treatments, as plots without flower access tended to have more adult ladybeetles than plots with flower access at the start of the study [Initial Adult Ladybeetle Density: BF: 3.7 ± 0.56 (mean ± SE); UBF: 2.5 ± 0.34 (mean ± SE); Two-Sample T-Test (Factor = Flower Access): t_14.9_ = 1.83, p = 0.087].

**Fig 2.**
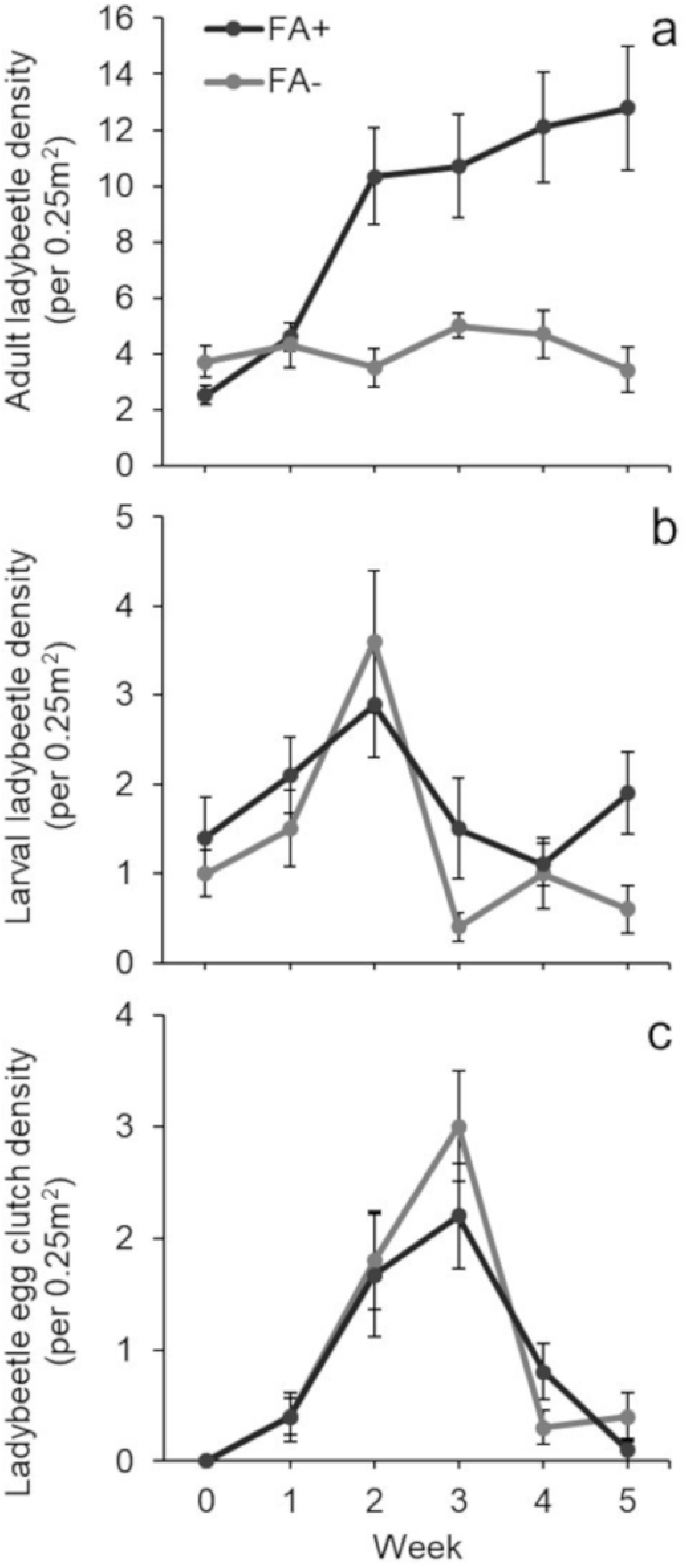
Effect of PRTs on ladybeetle population dynamics in the field. Mean (± SE) density of ladybeetle a) adults, b) larvae, and c) egg clutches in 0.25m^2^ manipulated field plots. Flower Access treatments (n= 10) are as follows: Flower Access Present (FA+) and Flower Access Absent (FA-).

Similar to effects on adult ladybeetles, larval ladybeetle density was impacted by Flower Access (F_1,107_ = 5.67, p = 0.019; S2 Table) and week (F_5,107_ = 5.08, p <0.001). Regardless of flower access, larval ladybeetle density peaked in all plots at week two (Fig 2b). However, the presence of cordgrass flowers increased larval ladybeetle density by 36% over the five-week study, while removing access to cordgrass flowers decreased larval ladybeetle density by 40% after five weeks.

The density of ladybeetle egg clutches depended upon time (F_5,107_ = 22.3, p <0.001; S3 Table), with clutch density peaking at week three in both treatments. Flower Access had no effect on egg clutch density (F_1,107_ = 0.38, p = 0.539; Fig 2c), despite adult ladybeetles being 4x more abundant in Flower Access Present plots.

Total scale insect density declined in both treatments over the five-week study (Fig 3). However, there was no difference between Flower Access treatments in the change in scale insect density during the study, despite the higher density of adult ladybeetles in plots with flower access (t_13.07_ = 0.347, p = 0.734).

**Fig 3.**
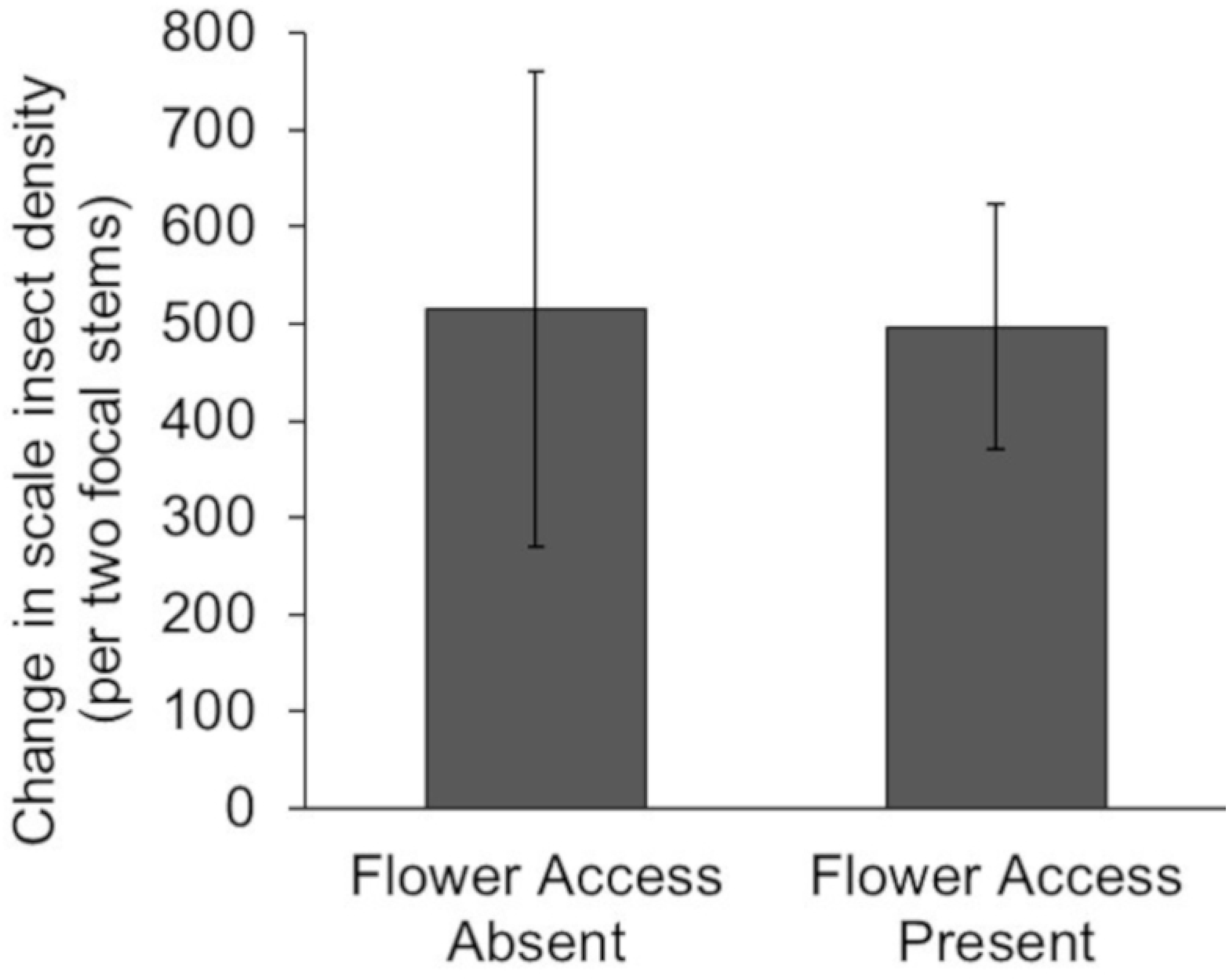
Effect of PRTs on ladybeetle foraging in the field. Mean (± SE) change in scale insect density on two focal cordgrass stems in our 0.25m^2^ manipulated field plots (n = 10).

### Effect of conspecific density on ladybeetle per capita scale insect consumption

In the presence of 1-3 conspecifics, adult ladybeetle density had no effect on the total number of scale insects consumed (linear regression: *R*^2^ = 0.07, p = 0.341; Fig 4a) and no cannibalistic activities between ladybeetles occurred. This result suggests that as conspecific density increased, per capita consumption of scale insects declined (linear regression: *R*^2^ = 0.489, p = 0.002 Fig. 4b). For example, adult ladybeetle per capita scale insect consumption (over three days) was 201 ± 58 (mean ± SE) for individual ladybeetles, but only 48 ± 20 (mean ± SE) for ladybeetles with two conspecifics (i.e. three adult ladybeetle treatment).

**Fig 4.**
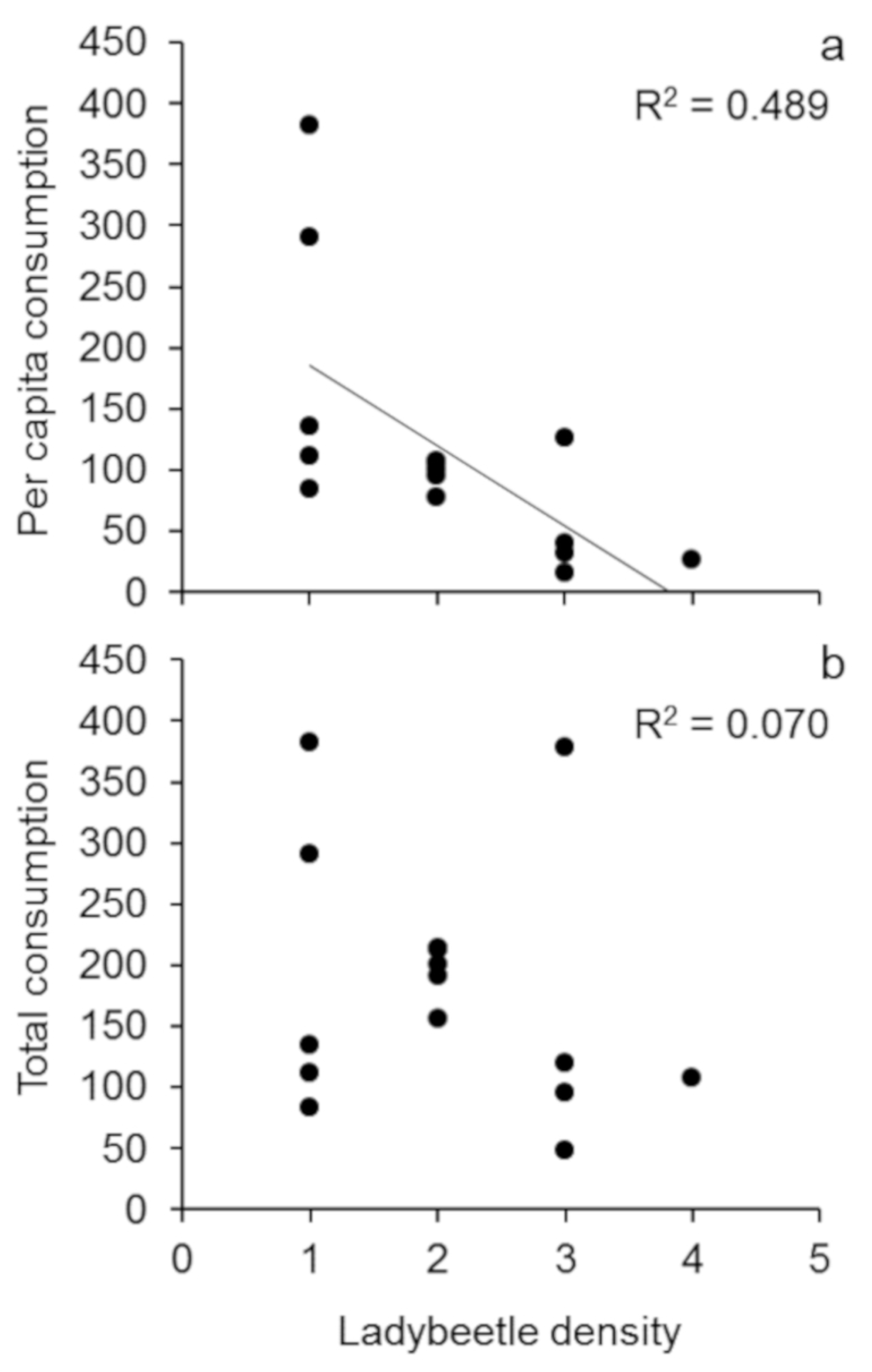
Effect of conspecific density on ladybeetle foraging rates. Mean (± SE) per capita consumption by adult ladybeetles of a) adult scale insects and **b)** total scale insects. Sample size five for all ladybeetle density treatments except the 3-ladybeetle treatment (n = 4) and 4-ladybeetle treatment (n= 1).

The number of ladybeetles observed on cordgrass plants (i.e., either leaves, stem, or flower) was not affected by Ladybeetle Density (F_2,12_ = 0.68, p = 0.523) or Observation Day (F_1,12_ = 4.00, p = 0.069; S4 Table, S1 Fig). For example, during the first observation day, replicates with 1 ladybeetle had 0.6 ± 0.24 (mean ± SE) ladybeetles present on the cordgrass plant, while replicates with 3+ ladybeetles had 1.0 ± 0.32 (mean ± SE) ladybeetles on the cordgrass plant. Similarly, on the second observation day replicates with 1 ladybeetle had 0.2 ± 0.2 (mean ± SE) ladybeetles present on the cordgrass plant, while replicates with 3+ ladybeetles had 0.4 ± 0.4 (mean ± SE) ladybeetles on the cordgrass plant. Thus, regardless of ladybeetle density within the replicate, ladybeetle density on the cordgrass plant appears to remain constant-suggesting that adult ladybeetles may avoid habitats already occupied by conspecifics.

## Discussion

Plant reproductive tissues (PRTs) can increase or decrease prey consumption by altering omnivore foraging behavior. In laboratory mesocosms, isolated adult ladybeetle prey consumption was unaffected by cordgrass flowers (Fig 1). In the field, habitat patches containing access to cordgrass flowers attracted 4x as many adult ladybeetles compared to habitats lacking flower access (Fig 2a). However, elevated ladybeetle densities in habitats with cordgrass flower access did not result in greater loss of scale insect prey (Fig 3), suggesting that PRTs resources reduced ladybeetle per capita consumption of scale insects. This discrepancy (PRTs had no effect in the lab but reduced per capita prey consumption in the field), may be related to intraspecific interactions (e.g., interference competition) between ladybeetles which were absent in the lab study with isolated ladybeetles. This hypothesis is supported by our finding that increasing conspecific density reduced per capita consumption of scale insects by ladybeetles. Overall, these observations suggest that PRTs may not impact the population level effects of omnivores on prey when numerical responses of omnivores are offset by reductions in their per capita predation rates.

PRTs decreased per capita consumption by omnivores on animal prey in our field study that allowed for intraspecific interactions between omnivores. Although access to PRTs (i.e. cordgrass flowers) increased adult and larval omnivore populations (412% and 36%, respectively), this did not translate into a change in animal prey density. Thus, our laboratory study of isolated adult omnivores and our field study contradict each other - access to PRTs reduced per capita consumption of animal prey by omnivores in the field but not the lab.

Omnivore feeding rates in the presence of PRTs may have differed in our laboratory and field experiments for two reasons. First, local environmental conditions may alter omnivore consumption of prey. For example, temperature can directly impact the metabolic rate of ectothermic omnivores, altering their energetic needs and, in turn, their foraging rates. However, we tried to minimize differences in environmental conditions between the laboratory and the field by 1) standardizing the month (August) of both experiments and 2) running our laboratory mesocosm study in outdoor seawater tables with natural tidal cycles (exposing experimental units to the ambient environmental conditions in southern California).

Second, the density of omnivores differed between our first laboratory (single omnivore) and field (multiple omnivores) studies. Differences in omnivore density may explain our conflicting findings if intraspecific interactions (e.g. mating, territoriality, or cannibalism) reduce omnivore consumption of focal prey resources. This seems likely, since our laboratory no-choice feeding assay found that conspecifics reduce ladybeetle per capita scale insect consumption (Fig 4). These findings parallel those of our field study, as ladybeetle populations in cordgrass flower habitats, despite being nearly 4x larger, removed the same number of scale insects as ladybeetles in plots lacking flower access (Figs 2 and 3). A recent meta-analysis suggests that the effects of PRTs on omnivore prey consumption depends on the ability of omnivores to elicit numerical responses. Specifically, in the presence of PRT, allowing omnivore numerical responses increased omnivore prey rate on animal prey, while not allowing numerical responses decreased omnivore predation rate on animal prey (Rinehart and Long *in prep.*).

While several studies have aimed to assess the impacts of PRTs on intraguild predation and cannibalism [see 9, 11-13], few have tested how elevated omnivore conspecific density (due to numerical responses to PRTs) may affect omnivore foraging behavior and local prey mortality. Here, we found that the presence of only two other conspecifics (i.e. 3+ ladybeetle treatment) decreased per capita prey consumption by 76% in just three days. Omnivores may consume fewer animal prey in the presence of conspecifics if they trade-off between foraging on prey and engaging in intraspecific interactions (e.g. mating or interference competition). For example, in our laboratory feeding assay, we frequently observed a single ladybeetle occupying a cordgrass plant at a time-regardless of ladybeetle density (i.e., 1,2, or 3+ ladybeetles). This observation suggests that adult ladybeetles may reduce their foraging efforts to avoid interacting with other ladybeetles at small spatial scales.

Changes in omnivore conspecific density and the availability of PRTs may also influence the rate of cannibalism between ladybeetles in our study system. For instance, post-aggregation cannibalism may explain why adult ladybeetle densities were 4x lower in field plots that denied ladybeetles access to flowers than those allowing ladybeetles access to flowers. However, cannibalism is unlikely in this system for two reasons. First, we never observed cannibalism between individual ladybeetles in the laboratory (including during our three-day no-choice assay) or the field (S.A. Rinehart *personal observation*). Second, cannibalistic events are most likely to occur when food resources, especially prey, are limited [12]. In all our studies, ladybeetles never consumed all scale insects in their environment, suggesting that the availability of prey was never limiting.

The effect of PRTs on prey consumption by omnivores is commonly attributed to nutritional benefits-as plants and animals vary in their nutrient, vitamin, mineral, and water content [41]. However, PRTs may also affect omnivore behavior by increasing habitat complexity. For example, habitat complexity can alter omnivore predation rates and antagonistic intraspecific interactions [42-43]. In our system, ladybeetles preferentially use cordgrass flowers as habitat - field surveys of randomly-selected cordgrass stems (n=95 individual flowering cordgrass stems) found that 88% of adult ladybeetles were found on cordgrass flowers versus other tissues (S5 Table).

The rate of omnivore prey consumption can be influenced by several factors. Historically, omnivory studies have focused on the impacts of PRTs on prey consumption and have found evidence that PRTs can both increase and decrease the rate of prey consumption by omnivores [5-7]. PRTs can increase local omnivore predation rates by attracting omnivores-as PRTs provide omnivores additional food resources and habitat structure [14-16]. However, few studies have tried to understand how local increases in omnivore conspecific density (due to aggregation to PRTs) ultimately affect omnivore-prey interactions. Here, we show that omnivore numerical responses to PRTs alter the foraging behaviors of omnivores, due to shifts in local conspecific density. Overall, our findings suggest a need to assess the indirect effects of PRTs on omnivore foraging behaviors to better understand how omnivory influences food web structure and function.

## Acknowledgements

We would like to thank B. Collins and S. Schroeter for access to Sweetwater Marsh and San Dieguito Lagoon. T. Grosholz, R. Karban, D. Deutschman, G. Vermeji, and J. Walker provided comments that improved the study design and final manuscript. F. Ventola, G. Cooper, and C. Knight provided field and laboratory assistance. Thank you to N. Fulner, K. Habersberger, Z. Kornfeld, and E.L. Yang for providing inspiration throughout this project. This is contribution No. XX of San Diego State University’s Coastal and Marine Institute. Author contribution statement: S.A.R conceived the project. S.A.R. and J.D.L. designed the study. S.A.R. performed the experiments and analyzed the data. S.A.R. and J.D.L. wrote the manuscript.

## Supporting information

**S1 Table. Repeated measures ANOVA for mean adult ladybeetle density between Flower Access treatments across the field six-week study.**

**S2 Table. Repeated Measures ANOVA for mean larval ladybeetle density between Flower Access treatments across the six-week field study.**

**S3 Table. Repeated Measures ANOVA for mean egg clutch density between Flower Access treatments across the six-week field study.**

**S4 Table. Repeated Measures ANOVA for number of ladybeetles on cordgrass plants between Ladybeetle Density treatments at two timepoints.**

**S1 Fig. Effect of conspecific density on adult ladybeetle behavior in laboratory no-choice feeding assays.** Mean (± SE) number of adult ladybeetles observed on cordgrass plant tissues in the laboratory no-choice feeding assay on the 2^nd^ and 3^rd^ days of the assay (n=5 per beetle density treatment).

